# Effect of urbanization and its environmental stressors on the intraspecific variation of flight functional traits in two bumblebee species

**DOI:** 10.1101/2021.01.29.428756

**Authors:** Nicola Tommasi, Emiliano Pioltelli, Paolo Biella, Massimo Labra, Maurizio Casiraghi, Andrea Galimberti

## Abstract

The way urbanization shapes the intraspecific variation of pollinator functional traits is little understood. However, this topic is relevant for investigating ecosystem services and pollinator health. Here, we studied how urbanization affects the functional traits of workers in two bumblebee species (*Bombus terrestris* and *B. pascuorum*) sampled in 37 sites along a gradient of urbanization in North Italy (an area of 1800 km^2^ including the metropolitan context of Milan and other surrounding capital districts). Namely, we investigated the role played by land-use composition, configuration, temperature, flower resource abundance, and air pollutants on the variation of traits related to flight performance (i.e., body size, wing shape and size fluctuating asymmetry). These traits are relevant because they are commonly used as indicators of stress during insect development. The functional traits of the two bumblebees responded idiosyncratically to urbanization. Urban temperatures were associated with smaller wing sizes in *B. pascuorum* and with more accentuated fluctuating asymmetry of wing size in *B. terrestris*. Moreover, flower abundance correlated with bigger wings in *B. terrestris* and with less asymmetric wing sizes in *B. pascuorum*. Other traits did not vary significantly and other urban variables played minor effects. These patterns highlight that environmental stressors linked to urbanization negatively impact traits related to flight performance and development stability of these species with possible consequences on the pollination service they provide.

Overall, this study found species-specific variation patterns in syntopic taxa, expanding our understanding about the effects of anthropic disturbance in shaping relevant functional traits of pollinator model species.

## INTRODUCTION

Widespread phenomena of urbanization are driving deep changes on landscape features, their temperatures and pollutants, creating novel ecosystem conditions that impact biodiversity (Foley et al., 2005; Weng et al., 2007; Wenzel et al., 2020). Plants and animals respond to these environmental variations by shifting their distribution (Colla et al., 2012), phenology (Huchler et al., 2020), and/or shaping some morphological traits considered “functional”, *i.e.* relevant for their ecology, fitness and behaviour (Alberti et al., 2017; Eggenberger et al., 2019; Nooten & Rehan, 2020). In bees, trait variation due to environmental alteration could affect the efficiency of the ecosystem services they provide though impacting the way they interact with plants (Buchholz & Egerer, 2020, Biella et al., 2019). Environmental alteration could also impact pollinator development, for example by limiting the abundance of floral resources due to the increasing proportion of anthropized surfaces (Steffan-Dewenter et al., 2001). This scenario, in turn, could trigger body size declines due to less food supplied to larvae (Couvillon et al., 2010). Furthermore, landscape anthropization could change the local climate, thus altering pollinator ecology, development and foraging (Radmacher & Strohm, 2010). Specifically, the higher degree of cemented “impervious” land cover that characterizes urban areas is often associated with increasing temperatures, a phenomenon known as the “heat island effect” (Chun & Guldmann, 2018). Observations from previous studies have strengthened the hypothesis that pollinator insects could face a shift towards smaller body size as an adaptation to reduce the risk of overheating while foraging in warmer conditions (Peters et al., 2016; Gérard et al., 2018a). Considering that the worldwide steady growth of the human population is driving a dramatic urban sprawl, new insights on pollinator responses are necessary.

Previous studies investigated the morphological responses of pollinators to anthropogenic pressures, mainly focusing on body size (e.g., Chown & Gaston, 2010; Eggenberger et al., 2019; Theodorou et al., 2021). In bees, this character responds rapidly to environmental changes (Chown & Gaston, 2010), it shows little heritability and its variation mainly depends on the amount of food received during the larval development (Couvillon et al., 2010). Bee size is positively correlated with the foraging range (Greenleaf et al., 2007). Generally, larger bees show more efficient flight performances (Harrison & Roberts, 2000), since flight muscle ratio is known to increase with body size in flying insects (Samejima & Tsubaki, 2010). Size also determines the metabolic rate and resource needs of adult imagos, with larger bees having higher metabolic rate (Kelemen et al., 2019) and thus potentially being more susceptible to shortage in floral resource availability (Couvillon & Dornhaus, 2010). However, to date, the investigation of pollinators body size variation in anthropogenic habitats yielded heterogeneous results. A recent study on bumblebees found bigger workers in cities (Theodorou et al., 2021). This study speculated that such pattern is an adaptation to longer flights for collecting resources, particularly in view of the severe green patches fragmentation of urban landscapes (Greenleaf et al., 2007). Conversely, a study by Eggenberger et al. (2019) found smaller bumblebee foragers in cities. This was interpreted as an effect of both limited local resource abundance and warmer temperature in urban areas, but the effect of these variables was not directly tested. Given these contradicting results and different interpretations, more studies are needed for clarifying the existing patterns of pollinator morphological responses to urbanization.

Morphometric studies are gaining in importance for quantifying even subtle variations in morphological traits. These variations are usually informative of stress exposure, and thus provide information about animal population health status (Adams et al., 2001). One of the advantages of using trait variation to measure stress is that changes of phenotypes are detectable before an overall decrease in population viability (Hoffmann et al., 2005). Therefore, quantifying traits variation could become an essential practice when evaluating local and landscape-level stressors. A metric that has grown in popularity is the fluctuating asymmetry (FA) (Klingenberg, 2001; Beasley et al., 2013; Alves-Silva et al., 2018), defined as the presence of small, randomly placed deviations from perfect bilateral symmetry due to the occurrence of developmental instability, driven by exogenous environmental conditions (Klingenberg, 2015). FA differs from another type of bilateral asymmetry, the directional asymmetry (DA), that occurs when the two sides are steadily different with a predictable direction to this difference. While DA has a genetic basis and therefore could be less impacted by the environment (Palmer & Strobeck, 2003), the FA is considered a valid proxy of stress exposure to conditions that typically occur in urban environments (e.g., higher temperature and air pollutants) (Beasley et al., 2013). For instance, laboratory-based studies have demonstrated that higher CO2 level or low temperature lead to an increase in wing FA, supporting the possible role of traffic pollutants and climatic variation in determining developmental instability (Klingenberg et al., 2001; Hoffmann, Collins & Woods., 2002). However, asymmetries could be found in wing shape and/or in wing size and they even have different responses to the same stressor type. For instance, in a recent study by Gerard et al., 2018, variations in wing size asymmetry were observed in response to thermic and parasitic stress while these same stressors caused no alteration in wing shape asymmetry level. Both wing size and shape are important functional traits in pollinators. This is because wing size is believed to be related to flight length and it influences metabolic costs (Fernandez et al., 2017; Soule et al., 2020), while shape is considered important for flight maneuverability (Kölliker et al., 2003, Grilli et al., 2017).

In order to characterize the effects of urbanization and of the related environmental stressors on pollinator insects, we quantified the morphological variation in two species of bumblebee *(i.e., Bombus pascuorum* and *B. terrestris).* We sampled foraging workers from populations spanned across a gradient of growing urbanization (from seminatural areas to highly urbanized sites) in Northern Italy, a region that experienced a strong anthropogenic footprint (Perini and Magliocco 2014; Salata 2017). The two species were selected as they are among the most common and widespread pollinators in Europe (Pekkarinen & Teräs, 1993; Rasmont et al., 2008), and have been largely used as model species in many studies related to the effects of urbanization or other stressors (Eggenberger et al., 2019; Theodorou et al., 2021).

We expected to find quantitative variation in bumblebee functional traits of body size and wing FA in response to several facets of urbanization. Firstly, we tested associations with increased fragmentation of green patches that is often found in urban landscapes (Li et al., 2019). We also tested responses due to environmental stressors amplified by urbanization, such as increased temperatures (Feng et al., 2014) and pollutants (Salahodjaev 2014), and decreased floral resource abundance (Ushimaru 2014). Regarding body size we based our survey on two alternative expectations that emerged from previous studies, depending on the prevalent pressure acting on this trait. First, one could expect to observe an increase in body size if green patches fragmentation triggered an adaptation to increase foraging ranges, as suggested by (Warzecha et al., 2016). Alternatively, a reduction in body size could arise as a way to reduce the risk of overheating in warmer urban habitats (Maebe et al., 2021; Pereboom & Biesmeijer, 2003) or as a consequence of limited floral resources (Chown & Gaston, 2010). Regarding wing FA in shape and size, we expected to find increased FA in response to higher levels of biotic and abiotic stressors that are expected to occur in more urbanized landscapes, such as limited floral resources, temperature, and air pollutants.

## MATERIALS AND METHODS

### Study species

This study was focused on two co-occurring species of bumblebee: *Bombus terrestris* (Linnaeus 1758) and *B. pascuorum* (Scopoli 1763). Both species are pollinators common in Europe, and can be easily found while foraging in different habitats (Polce et al., 2018), even in urban areas (Meeus et al. 2021, Banaszak-Cibicka, and Żmihorski 2012), including the surveyed region (personal observation of the authors). Given these characteristics, these species are reliable models to investigate responses by pollinating insects to landscape anthropization (Eggenberger et al., 2019; Theodorou et al., 2021). Using two cases of different, albeit related, model species could even clarify if the observed patterns are general or rather shaped by different life histories. The two selected species, in fact, have slightly different foraging ranges, with an estimated maximum of 449 and 758 m for *B. pascuorum* and *B. terrestris,* respectively (Knight et al., 2005). Nesting habits are also dissimilar as *B. terrestris* builds its nest in subterranean holes, while *B. pascuorum* on top of or slightly beneath the soil surface (Goulson, 2010). Another important difference is represented by their dietary regimes, since *B. pascuorum* usually have a narrower trophic niche and a preference for deep-corolla flowers (Harder, 1985) while *B. terrestris* is highly polylectic (Dafni et al., 2010, Biella et al., 2019).

### Study design and sampling

Samplings were conducted at 37 sites (Fig. 1), in July 2019, between 9:00 and 12:00 only on days with sunny and windless weather conditions. The study sites were distributed within an area of about 1800 km^2^ covering four administrative provinces, Milano, Monza e della Brianza, Lecco and Como in northern Italy. A minimum distance between sites of 1 km was imposed to avoid the non-independence of sites (Phillips et al., 2019) since it is above the maximum foraging range observed for the two species (Knight et al., 2005). Study sites were selected along a gradient of growing urbanization, ranging from areas highly dominated by seminatural hay meadows close to forest with little urban areas nearby, to sites characterized by a high degree of impervious surface (i.e., concrete, building, and asphalt). To select sampling sites, impervious surfaces were mapped in a GIS software based on a regional land use cartography (2018-DUSAF 6.0; https://www.dati.lombardia.it/Territorio/Dusaf-6-0-Uso-del-suolo-2018/7rae-fng6) This land use cover map is available at the scale of 1: 10000 and was developed from AGEA orthophotos and SPOT 6/7 satellite images.. Sites were chosen on a visible gradient of growing impervious cover. Once identified as suitable, the land use composition in the surrounding of the candidate sampling sites was confirmed using satellite images. For each species, five to six worker specimens were captured while foraging inside a plot of about 50 m x 50 m at each site using an entomological net. After collection, the insects were stored at −80 °C until further analyses.

**Fig.1:**
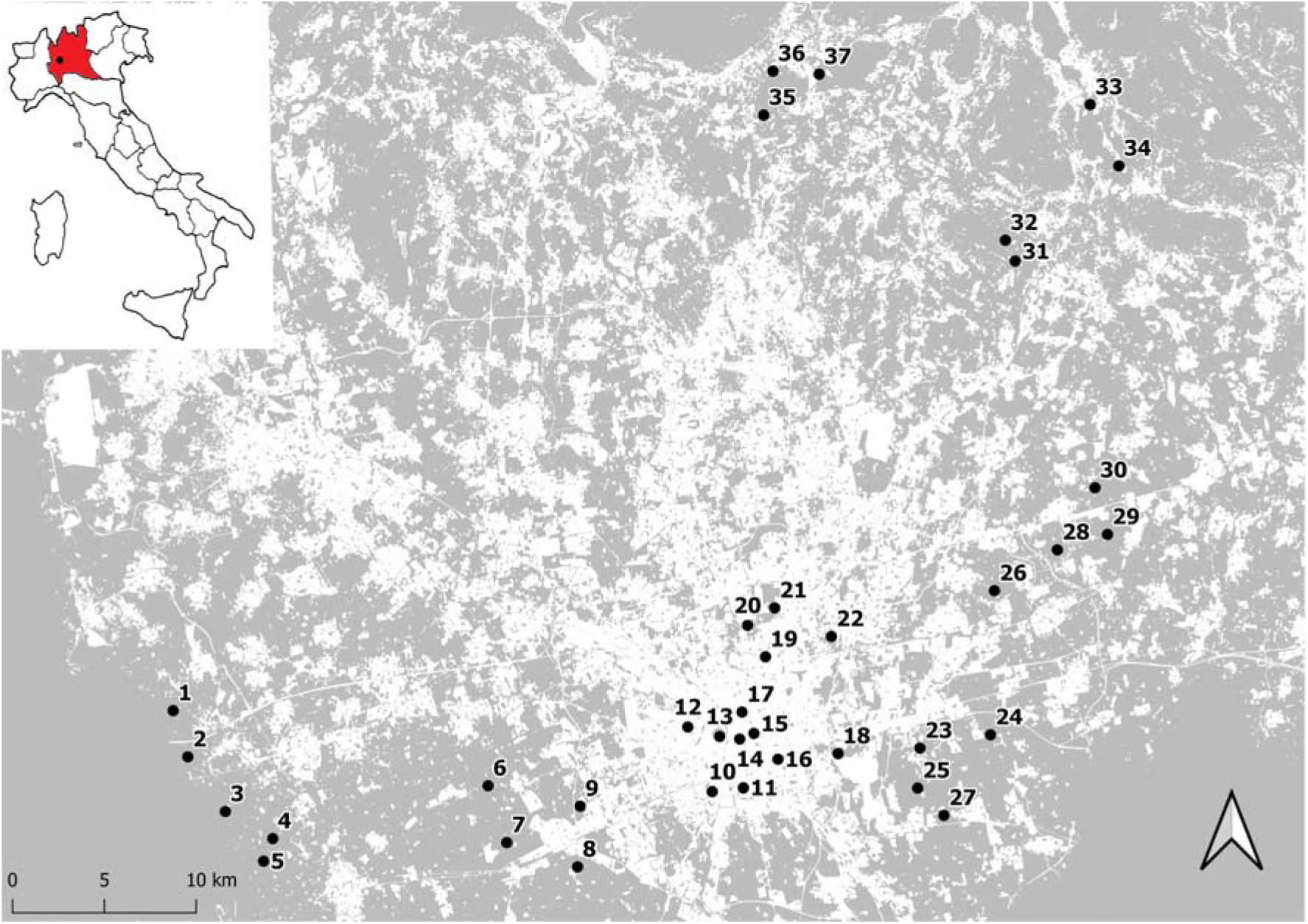
Map showing the distribution of the sampling sites along the urbanization gradient. White areas correspond to cemented surfaces.

### Landscape and environmental variables description

The previously mentioned land use cartography was used to quantify landscape urbanization around sampling sites. Through QGIS 3.10.11, a 1 km radius buffer area was created around each site where landscape composition was evaluated arranging DUSAF original level and sub-level of land-use classification into two macro categories: impervious (i.e., buildings, infrastructures, roads, and cemented surface), and seminatural land (i.e., meadows, forest and urban green spaces) (see Online resources, Additional information 2 for a list of DUSAF codes assigned to each grouping).

For each site the ratio between impervious and green land was computed to quantify the intensity of urbanization. The gradient of urbanization was also described by green habitat fragmentation, a measure of landscape configuration, that was quantified by computing the edge density (ED), as the ratio of edge length of green and seminatural patches over their total area (Wang et al., 2014). Other environmental biotic and abiotic features, possibly influenced by the urbanization degree, were considered to test for their potential effects on altering body size and wing size/shape FA. Specifically, land surface temperature was calculated as the mean value in the period June-July using data retrieved through remote sensing imaging spectroradiometer (MODIS) MOD11A2 from the NASA database (https://modis.gsfc.nasa.gov/data/dataprod/mod11.php) with a resolution of 1 km. As the two species of bumblebee studied are characterized by a life-cycle from eggs to adults of about two months (Goulson, 2010), the time frame for which these data were taken into consideration should well represent the mean temperature experienced by larvae during their development. It is important to notice that at the resolution used here it is not possible to properly describe microclimatic variation, but rather broader temperature variation covering the landscape scale and the foraging range of the two bumblebee species (Knight et al., 2005). A map reporting the variation of mean temperatures along the investigated landscape is reported in Online resources, Figure S1. Air pollution was estimated as the mean of daily concentrations of NO_2_ over two months (June and July) registered by Regional agency for environmental protection (ARPA), data taken from monitoring stations located nearby our sampling sites (https://www.arpalombardia.it/Pages/Aria/qualita-aria.aspx). A map reporting the location of monitoring stations along the investigated landscape is provided in Online resources, Figure S1.

An expeditive estimation of floral resources at each site (i.e., the total number of flowers) was performed by using six quadrats 1 m x 1 m (covering a proportion of sampling area similar to that reported in Fisher et al., 2017) randomly placed in the flowering green spaces within or closest to the sampling area, and counted the number of flowers found there (as in Ushimaru 2014). Flowers were counted considering single or composed inflorescences as units: for *Myosotis* sp., *Galium* sp., and *Capsella bursa-pastoris*, and all Asteraceae the number of inflorescences was counted. The values of the listed landscape and environmental variables in all the sampling sites are reported in Online resources, Table S1 along with histograms showing their variation along the sites (Additional information 1)..

### Specimens imaging and wings measurement

The forewings of all individuals were detached at the base and scanned at high resolution (i.e., 600 dpi). The obtained images were converted into TPS files using tps-UTIL 1.74. This file format follows the standard formats for geometric morphometrics (Rohlf, 2015). TPS file can contain two or three dimensional landmark data and the information about the scale factor applied to each specimen is also provided. Once converted into TPS, images were digitised using the tps-Dig 2.31 software (Rohlf, 2015), with two-dimensional cartesian coordinates of 15 landmarks positioned at wing vein junction (Fig. 2) (as in Aytekin et al., 2007; Klingenberg et al., 2001). Bumblebees with damaged or badly worn wings were excluded from further analyses.

**Fig.2:**
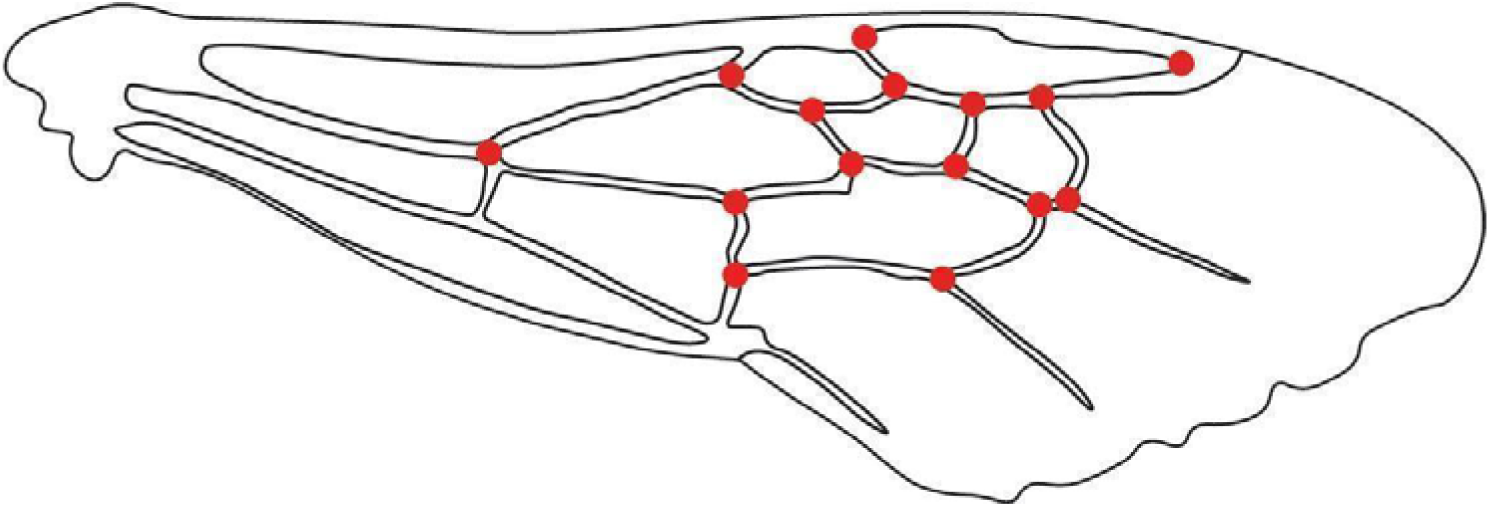
Right forewing of *B. terrestris* with landmark locations used in this study. Details on the formulas applied to calculate centroid size and consequently fluctuating asymmetry from these landmarks are reported in the manuscript section “Specimens imaging and wings measurement” and in the references within.

The analysis of landmark configuration was conducted in MorphoJ 1.07 software (Klingenberg, 2011). To remove all the effects of scale, rotation and position, a standard protocol based on a generalized least square Procrustes superimposition was applied (Klingenberg, 2011). This strategy permits to obtain a new set of superimposed landmark coordinates (i.e., ‘Procrustes shape coordinates’) describing the wing shape and size features. Wing size was estimated as the centroid size: *i.e.*, the square root of the sum of squared distances from the centroid of each landmark configuration, and used as a proxy of body size (hereafter “body size”, as in Outomuro & Johansson, 2011 and Dellicour et al., 2017). To confirm the positive relation between centroid size and body size the inter tegular distance (IT), another measure of body size (Warzecha et al., 2016), was retrieved from a subset of 50 individuals of each species. Afterwards, the correlation between IT and centroid size was calculated (*B. terrestris* r = 0.7, p <0.001; *B. pascuorum* r = 0.7, p <0.001). Wing size asymmetry was computed by dividing the absolute difference between left and right centroid sizes by the mean centroid size and multiplying by 100 (Leonard et al., 2018). To estimate wing shape variation, Procrustes distances were computed for each individual (Klingenberg, 2015). These represent the measure of an individual’s overall asymmetry (*i.e.*, combining DA and FA components), obtained by taking the square root of the sum of squared distances between corresponding right and left Procrustes’ coordinates (Klingenberg, 2015).

### Statistical analysis

According to the protocol outlined in Klingenberg 2015, we first estimated the entity of the measurement error because the levels of asymmetry in bilateral traits are subtle and it could possibly cause considerable variation in the assessment of asymmetry levels. This was performed by double-scanning the wings and digitizing their landmarks for a subset of 40 specimens, with the Procrustes ANOVA in MorphoJ (Klingenberg 2001, Klingenberg 2015) in order to evaluate the measurement error. Afterwards, following Costa et al. (2015), to isolate the FA component, we calculated the amount of directional asymmetry (DA) and tested its entity again with the Procrustes ANOVA in MorphoJ considering all the measured specimens in a single analysis. Only if DA occurred significantly, the individual asymmetry measures were corrected by subtracting the mean DA, thus isolating the FA component as in Costa et al. (2015).

To investigate the relationship between morphological traits and covariates, linear mixed models were used. The responses of the two species were assessed separately. In all the models, the ratio between impervious and seminatural surfaces was initially included as a predictor with the other variables. However, Variance inflation factor (VIF) criteria was used to assess the absence of collinearity among model variables and it indicated that the ratio between impervious and seminatural surfaces was highly collinear with the other variables, see also the correlation matrix reported in Online resources, Table S2. Thus, we decided to exclude the ratio between impervious and seminatural surfaces from subsequent models. The other variables, describing landscape configuration, biotic, and abiotic features, as well as the interaction between all these variables, were included in the models following the ecological expectations of our hypothesis. Specifically, changes in body size were evaluated in response to the edge density of green area, temperature, and floral resource availability because they could directly influence body size, with bigger sizes in increasingly fragmented green areas, and/or with more flower resources, and/or less temperatures (Warzecha et al., 2016; Pereboom & Biesmeijer, 2003; Chown & Gaston, 2010). Concerning wing FA, the temperature, concentration of NO_2_, and flower resources limitation were included in the models following our hypothesis that they could be stressors expected to increase asymmetry (Hoffmann, Collins & Woods, 2002; Klingenberg et al., 2001; Leonard et al.,2018) Sampling site was included as a random effect in all the models. A backward stepwise model selection based on AIC was used to remove variables and their combinations that did not improve the model fit and thus to obtain reliable final models (Zuur et al., 2009). In order to improve the fit between the predictors and the response variable, mathematical transformations were applied on some of the covariates as reported in Table 1. All the analyses were performed using R (version 3.6.1; R CoreTeam 2019).

**Table 1:** Output of Linear mixed models of body size (N= 348) and fluctuating size asymmetry (size FA) (N= 347) of each species as a function of biotic and abiotic covariates of urbanization, with site identity as random factor. Final models were selected through backward stepwise selection using AIC criterion. ΔAIC reports the difference in AIC values between full and final models. β_i_: regression coefficient; χ^2^: chi square values; df: degrees of freedom. Models and results of shape FA are reported in Online resources, Table S3 as they were non significant.

| Species | Response variable | Full model covariates | Final model covariates | $\Delta AIC$ | $\beta_i$ | $\chi^2$; df | p value |
| --- | --- | --- | --- | --- | --- | --- | --- |
| *Bombus terrestris* | Body size | Temperature T<br>Edge density ED<br>log (Floral resources FL)<br>Interaction T x ED x FL | log (Floral resources) | 16.5 | 0.025 | 6.610;1 | **0.010** |
| *B. pascuorum* | Body size | Temperature T<br>Edge density ED<br>log (Floral resources FL)<br>Interaction T x ED x FL | Temperature | 17.9 | -0.003 | 7.403;1 | **0.006** |
| *B. terrestris* | Size FA | Temperature T<br>log (Floral resources FL)<br>NO<sub>2</sub> N<br>Interaction Tx FL x N | Temperature | 23.5 | 0.052 | 7.183 | **0.007** |
| *B. pascuorum* | Size FA | Temperature T<br>log (Floral resources FL)<br>NO<sub>2</sub> N<br>Interaction Tx FL x N | log (Floral resources) | 30.3 | -0.161 | 6.118;1 | **0.013** |

## RESULTS

After excluding queen, males, and specimens presenting damaged wings, 179 *B. pascuorum* (mean per site = 4.8 ± 0.4) and 169 *B. terrestris* (mean per site = 4.5 ± 0.3) were subjected to morphometric analyses.

The measurement error was negligible because it was not significant for wing size (df = 79, F = 2.67 p = 0.4578, R^2^ = 0.0009) and shape (df = 2054, F = 0.51, p = 0.9976, R^2^ = 0.07). Different patterns of size variation were found in the two bumblebee species. *B. terrestris* body size was found to increase in response to floral resource abundance (Fig. 3a, Table 1) while *B. pascuorum* body size decreased in response to the increasing temperature (Fig. 3 b, Table 1).

**Fig. 3:**
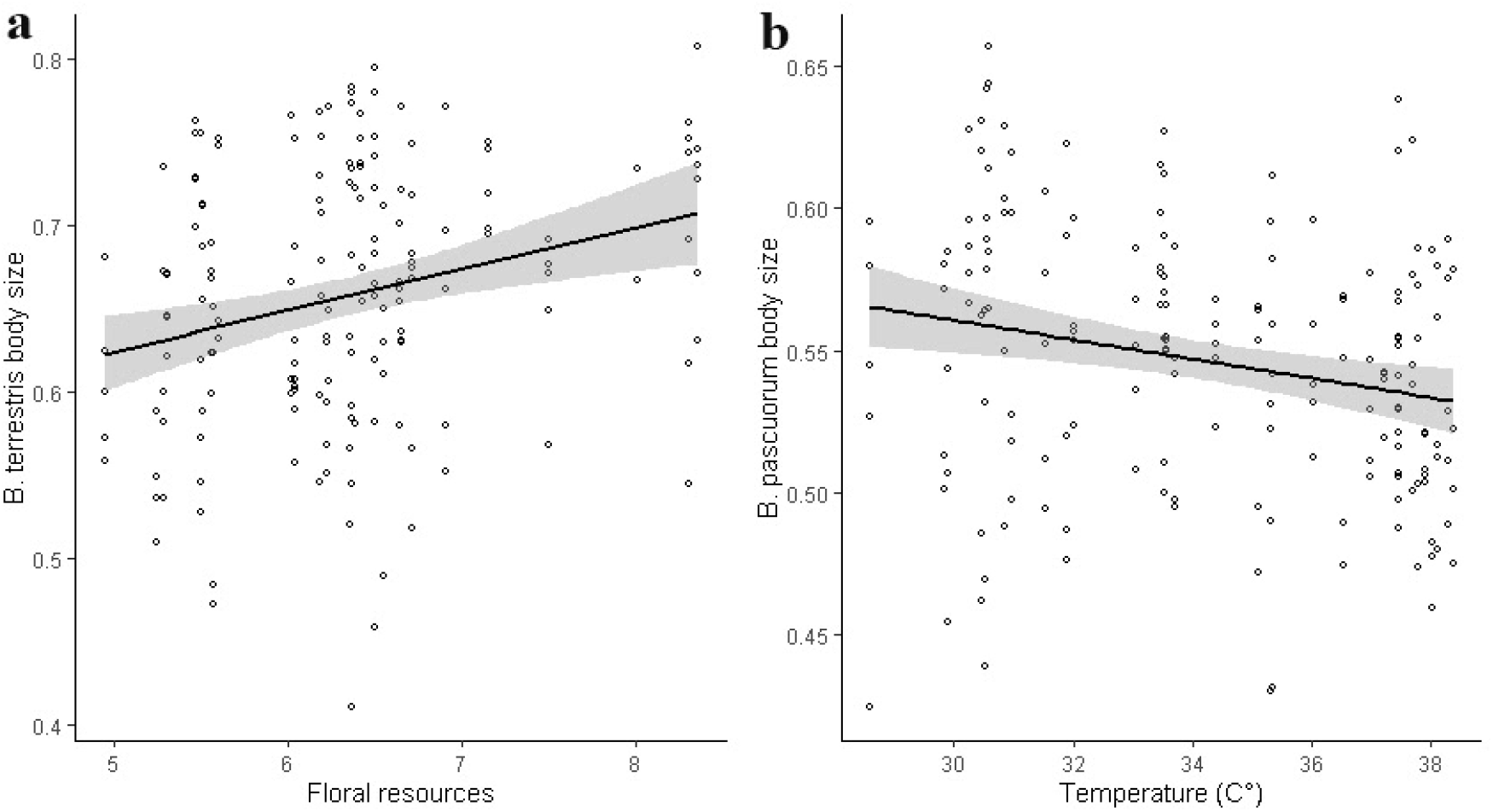
Body size variations as a function of (a) Floral resource abundance in *B.terrestris* and (b) Summer temperature in *B. pascuorum*. The black line and grey areas indicate the relationship and its confidence intervals as estimated with Linear mixed models, see methods for further details.

Concerning wing asymmetry, both species showed a significant level of shape DA (*B. pascuorum* df = 26, F = 4.66; p < 0.0001, R^2^ = 0.008; *B. terrestris* df = 26, F = 5.60; p < 0.0001, R^2^ = 0.009), while size DA was statistically significant only in *B. pascuorum* (df = 1, F = 29.77; p < 0.0001, R^2^ = 0.0007; in *B. terrestris* df = 1, F = 0.51; p = 0.4779, R^2^ = 0.000008). Measures of FA were then obtained by correcting for directional components.

*B. terrestris* size FA was positively correlated to temperature (Fig. 4 a; Table 1) while *B. pascuorum* size FA decreased with higher floral resource availability (Fig 4 b; Table 1). None of the predictor variables (i.e. temperature, NO_2_, resource abundance, and the interaction among these variables) showed a significant effect on wing shape asymmetry in both bumblebee species (output of non significant regression models are available in Online resources, Table S3).

**Fig. 4:**
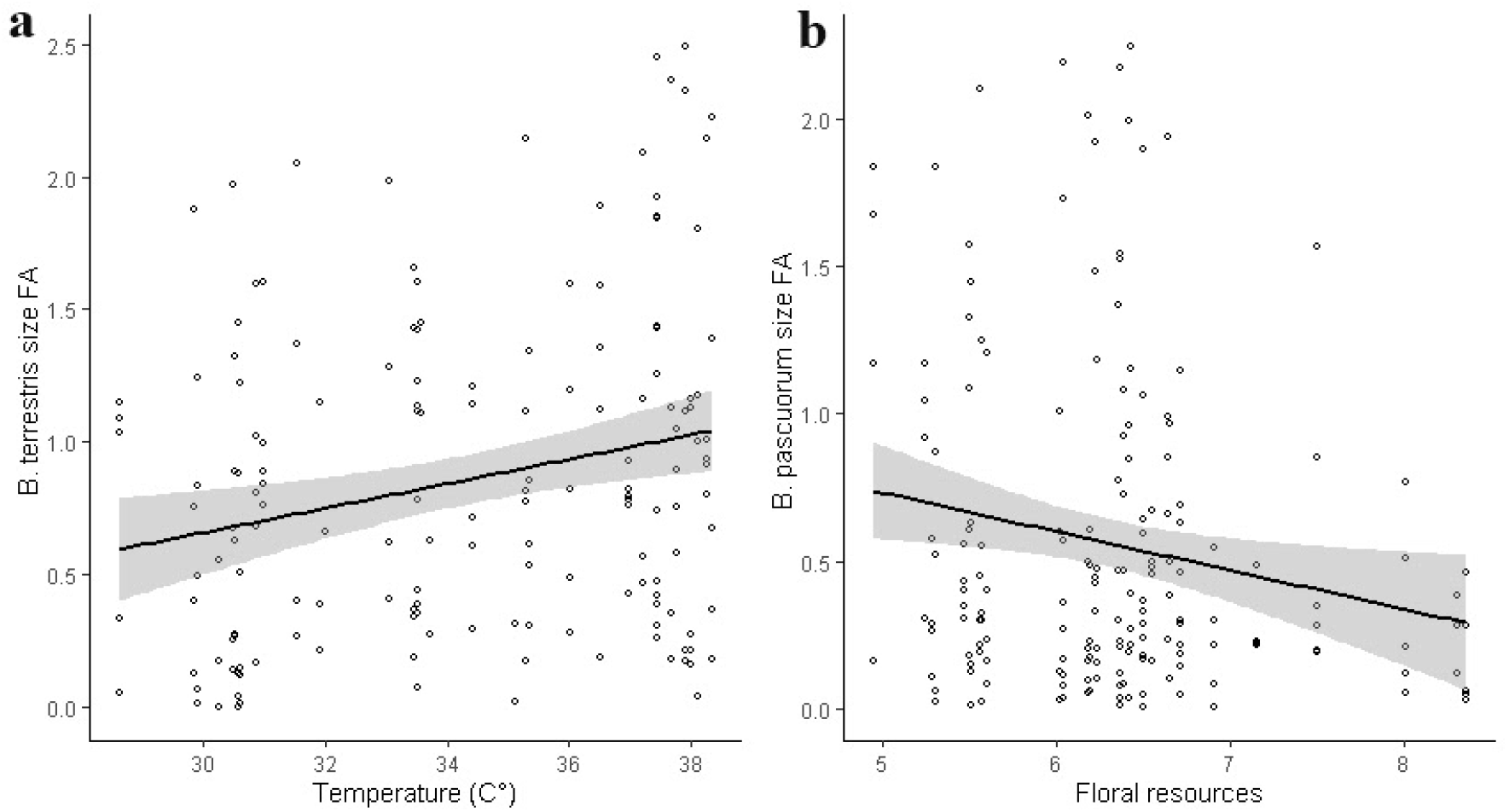
Variations in wing size Fluctuating Asymmetry (FA) as a function of (a) Summer temperature in *B. terrestris* and (b) Floral resource abundance in *B. pascuorum*. The black line and grey areas indicate the relationship and its confidence intervals as estimated with Linear mixed models, see methods for further details.

## DISCUSSION

In this study, we quantified the spatial intraspecific variation of functional traits in two common bumblebee species, to infer the possible alteration of their ecological features, such as the dispersion ability, with consequences on different aspects of their life history.

We focused on the morphological variations of *B. terrestris* and *B. pascuorum* along an urbanization gradient within an area of 1800 km^2^ including the metropolitan context of Milan and other surrounding capital districts. Our results highlighted some correlations between stressors related to urbanization, and traits as body size and wing size asymmetry. Specifically, the landscape temperature and the abundance of floral resources, two environmental features influenced by the degree of urbanization (See online resources Table S1, Additional information 1, and Table S2), emerged as candidate drivers of intraspecific variation of body size across bumblebee populations, acting differently on the two investigated species. Foragers of *B. pascuorum* showed a shift towards smaller body size in response to increasing temperature, a condition often associated with deeply urbanized landscapes, generally referred to as the heat island effect. Although a similar pattern of body size reduction in urban bumblebees has previously been reported by Eggenberger et al. (2019), they did not evaluate the effect of temperature but only proposed it as a possible driver of the observed decrease of body size. This relationship has been investigated on multiple historical series collections-based and experimental studies that revealed how higher environmental temperatures represent a driver of body size reduction in different species of bees (e.g., Nooten & Rehan, 2020; Theodorou et al., 2021). Higher temperature accelerates larval development, which likely results in smaller adults (Sibly & Atkinson, 1994). Furthermore, smaller sizes in warmer areas could also be a strategy for reducing overheating risks while foraging, due to an increased convective heat loss in smaller bees (de Farias-Silva & Freitas, 2020). Functionally, smaller foragers could travel shorter foraging distances (Greenleaf et al., 2007) and could also load less pollen and nectar (Goulson et al., 2002). As a consequence, the shift towards smaller body size in *B. pascuorum* could imply that it will pollinate less plants or handle flowers less efficiently (Földesi et al., 2020), a concerning aspect in view of colony provision and pollination. It is not known if microclimatic conditions could mitigate the effects we observed, as we used temperature measured at a broader scale. Furthermore, it is important to underline that other landscape variables could act synergically with urban temperature. However, these synergic effects could be difficult to disentangle in the field, due to the high correlation of these variables with the amount of impervious land. Further investigations considering microclimatic variations (e.g., by using data loggers at each sampling site) and field experiments pointing at cause-effect relationship between temperature and body size will be required to exclude the possible role of other urban related stressors. The role of different categories of impervious surface (i.e., concrete, building, and asphalt) in contributing to temperature increase should also be addressed in future research, to better inform mitigation strategies in urban contexts.

Body size reduction was also previously explained by the decrease in floral resource abundance in urban landscapes, possibly as a consequence of reduction of green areas (Merckx et al., 2018; Eggenberger et al., 2019). According to this evidence, we found a correlation between *B. terrestris* body size and the abundance of flower resources, with larger individuals observed where more food is potentially available. This is in accordance with the observation that adult size is strictly correlated with the amount of food received during larval development (Couvillon & Dornhaus 2009). However, this trend seems to be not clearly confirmed by *B. pascuorum* probably due to a possibly higher flower specialization of this species (Harder, 1985). Indeed the narrower diet of *B. pascuorum* could result in the inability of this species in exploiting abundant flowering plant species that do not represent its host plants. It seems a fruitful avenue of future research to investigate other important features such as the nutritional quality of available resources, their diversity, and changes along landscape variation (Vaudo et al., 2015; Vaudo et al., 2016). Here, we considered floral resource abundance at the local scale. In spite of this limitation, we could observe some relationships between traits and flower resources. However, additional data will further illuminate how resources shape urban pollinator traits.

Despite *B. pascuorum* and *B. terrestris* belong to the same genus, they showed a different susceptibility towards the investigated stressors. This suggests that different responses are likely to come from different habits and behaviour. Idiosyncratic responses were also observed in other bumblebee species, where body size decreased over warming decades, but others responded in the opposite way (Gérard et al., 2020). In our study the invariant size of *B. terrestris* in warmer conditions could be explained by its higher heat tolerance (Martinet et al., 2020). In addition, *B. terrestris* nests further underground compared to *B. pascuorum*, and it might be less exposed to warm air temperatures during larval development. These aspects strengthen the hypothesis that temperature could be a major determinant of pollinator size reduction in cities because they affected the body size of the more temperature-sensitive species, but not the heat-tolerant one. These idiosyncratic species-specific responses are very relevant for understanding the potential mechanism of intraspecific trait variation associated with urbanization and supports the need to consider a wider panel of species in this kind of studies.

Regarding wing asymmetry we found that size FA was positively correlated with increased temperatures in *B. terrestris*. Variation in both wing size and wing shape asymmetry was observed in other insect taxa and the effect of temperature was previously investigated under controlled laboratory conditions (Mpho et al., 2002). Studies associated the increased wing size and shape FA to environmental stressors, indicating that impairment of developmental processes might take place (e.g., Klingenberg et al., 2001; Kerr et al., 2013). The absence of variation in shape asymmetry registered for both the bumblebee species could confirm the results from other studies that have indicated shape variation as less susceptible to stressors than size asymmetry (e.g., Gérard et al., 2018b). Importantly, floral diet could represent a possible mitigation of environmental stressors during development (Archer et al., 2014). Here, this view is supported by the negative correlation found between resource abundance and wing size FA, although only in *B. pascuorum*.

Flight performance largely depends on body size, and is also affected by asymmetries in shape and size between wings (Grilli et al., 2017; Soule et al., 2020). Variation in these traits does not only show developmental instability, but also has deep ecological implications. Indeed, body size is determinant in predicting dispersal ability of insects, thus determining their foraging range (Greenleaf et al. 2007). Similarly, wing size FA impacts the management of lengthy flights (Fernandez et al., 2017, Soule et al., 2020), while wing shape FA is often associated with flight maneuverability. The combination of these morphological changes could deeply impact flight performance, flight length in time and space, and consequently bumblebee foraging (Kenna, Pawar & Gill., 2021), with potential consequences on their pollination efficiency. However, an important aspect to consider is that wing size and its asymmetry could even determine behavioural changes. For example, insects could increase visitation rates at closer distances to colonies, and even spend a higher time on the available resources instead of flying at a broader distance (Andrieu et al., 2009). Such changes, albeit difficult to quantify in the field, could merit further investigation when trying to forecast the impact of functional trait changes in response to urbanization.

## CONCLUSIONS

This study suggests that the environmental changes associated with urbanization could affect different functional traits of pollinators, and that their impact occurs heterogeneously on different insect species. Eventually, as the studied traits are often involved in flying abilities, these responses could bring to the alarming outcome of decreased foraging efficiency and pollination effectiveness. Furthermore, the different responses to the same stressor among the two bumblebees underline the necessity for future studies to consider a wider panel of taxa instead of single model species in order to draw conclusions that could be applied to the whole pollinator insects community.

From a conservation perspective, the comprehension of how pollinators cope with the challenging conditions occurring in novel anthropogenic habitats, plays a key role in informing suitable policy efforts to conserve their biodiversity and the ecosystem service they provide. In the future, cities are predicted to expand constantly and thus designing of urban landscapes will become a fundamental step for achieving sustainability outcomes. The pollinator-friendly design and management of urban green spaces will possibly create suitable conditions for pollinators and thus for the ecosystem services they provide (Guenat et al., 2019; Tommasi et al., 2021). At the same time, urban forestry and greenery practices (e.g., plantation of street and residential trees and the creation of urban greenbelts or greenways) could represent a valid solution to mitigate stressful conditions related to the urban environment, such as the lack of floral resources and the heat island effect (Chun & Guldmann, 2018) that here were found to influence functional traits.

## Supporting information

online resource

## ACKNOWLEDGMENTS

The authors are grateful to Amelia Pioltelli for artworks, Giulia Masoero, Lorenzo Guzzetti, Carota, Sam and Jemy for useful advice and support during manuscript preparation.

## DECLARATIONS

### Ethics approval

Sampling permits were obtained when needed from local authorities

### Availability of data and material

All relevant data are within the paper or stored in a public repository (https://doi.org/10.6084/m9.figshare.13637594).

