## Supplementary material for "Effect of urbanization and its environmental stressors on the intraspecific variation of flight functional traits in two bumblebee species": online resource

### Table of contents

**Table S1 -** Table of variables distribution among sampling sites – **pag 2**

**Additional information 1 -**Histograms representing variables distribution among sampling sites – **pag 3-5**

**Table S2 -** Correlation matrix between variables – **pag 5**

**Figure S1 -** Map of mean temperature and $NO_2$ sampling points **- pag 6**

**Table S3 -** Complete regression models outputs – **pag 7**

**Additional information 2 -** Description of DUSAF levels categorised as “Impervious” and “Semi Natural” – **pag 8**

### TABLE S1 - TABLE OF VARIABLES DISTRIBUTION AMONG SAMPLING SITES

| SITE CODE | Impervious/semi natural surfaces | Edge density | Min C° | Average C° | Max C° | Min NO2 µg/m3 | Average NO2 µg/m3 | Max NO2 µg/m3 | Floral resources |
|---|---|---|---|---|---|---|---|---|---|
| 1 | 0.0 | 0.040 | 28.6 | 30.3 | 32.7 | 7.1 | 22.44 | 41.100 | 510 |
| 2 | 0.1 | 0.045 | 28.4 | 30.6 | 33.2 | 10.4 | 26.87 | 49.800 | 823 |
| 3 | 0.0 | 0.020 | 27.2 | 29.9 | 32.8 | 2.0 | 15.25 | 32.800 | 198 |
| 4 | 0.1 | 0.055 | 28.6 | 30.5 | 32.2 | 2.0 | 15.25 | 32.800 | 1001 |
| 5 | 0.0 | 0.043 | 27.7 | 29.9 | 31.9 | 2.0 | 15.25 | 32.800 | 141 |
| 6 | 0.0 | 0.069 | 30.9 | 33.1 | 35.5 | 8.0 | 18.17 | 42.000 | 490 |
| 7 | 1.9 | 0.088 | 29.6 | 32.0 | 33.8 | 10.7 | 23.82 | 51.900 | 622 |
| 8 | 1.2 | 0.069 | 31.4 | 33.5 | 35.5 | 10.7 | 23.82 | 51.900 | 4049 |
| 9 | 1.5 | 0.077 | 32.8 | 35.1 | 37.5 | 19.0 | 26.12 | 31.000 | 3020 |
| 10 | 8.6 | 0.093 | 36.2 | 38.3 | 40.7 | 16.1 | 30.77 | 52.400 | 662 |
| 11 | 8.5 | 0.063 | 36.1 | 38.1 | 40.5 | 16.1 | 30.77 | 52.400 | 190 |
| 12 | 11.6 | 0.073 | 34.9 | 37.2 | 39.5 | 16.1 | 30.77 | 52.400 | 583 |
| 13 | 5.3 | 0.038 | 35.2 | 37.5 | 40.1 | 16.1 | 30.77 | 52.400 | 583 |
| 14 | 6.7 | 0.051 | 35.2 | 37.7 | 40.3 | 16.1 | 30.77 | 52.400 | 270 |
| 15 | 8.1 | 0.053 | 35.8 | 37.9 | 40.5 | 16.1 | 30.77 | 52.400 | 244 |
| 16 | 14.1 | 0.065 | 36.3 | 38.0 | 39.9 | 16.1 | 30.77 | 52.400 | 577 |
| 17 | 17.4 | 0.081 | 35.6 | 38.4 | 40.8 | 16.1 | 30.77 | 52.400 | 410 |
| 18 | 2.3 | 0.044 | 35.2 | 37.5 | 39.3 | 16.1 | 30.77 | 52.400 | 1273 |
| 19 | 11.8 | 0.083 | 36.1 | 37.8 | 39.5 | 21.3 | 40.89 | 57.600 | 826 |
| 19 | 0.2 | 0.071 | 31.4 | 33.6 | 34.9 | 10.0 | 19.29 | 38.000 | 420 |
| 20 | 1.5 | 0.046 | 34.9 | 36.5 | 37.9 | 21.6 | 39.94 | 53.500 | 248 |
| 21 | 1.5 | 0.023 | 34.2 | 36.0 | 38.0 | 21.3 | 40.89 | 57.600 | 201 |
| 22 | 2.9 | 0.086 | 35.3 | 37.4 | 39.5 | 31.0 | 38.19 | 55.000 | 1805 |
| 23 | 1.3 | 0.069 | 33.9 | 35.4 | 36.9 | 12.0 | 17.62 | 29.000 | 420 |
| 24 | 1.3 | 0.057 | 31.7 | 33.5 | 35.1 | 12.0 | 17.62 | 29.000 | 664 |
| 25 | 0.4 | 0.096 | 33.1 | 35.3 | 36.8 | 12.0 | 17.62 | 29.000 | 612 |
| 26 | 0.5 | 0.036 | 31.5 | 33.7 | 35.6 | 15.0 | 22.69 | 28.000 | 592 |
| 27 | 1.4 | 0.069 | 33.9 | 37.0 | 39.0 | 14.0 | 17.5 | 25.000 | 4270 |
| 28 | 1.1 | 0.085 | 32.5 | 34.4 | 35.8 | 19.1 | 32.06 | 49.700 | 765 |
| 29 | 0.1 | 0.083 | 31.6 | 33.5 | 34.8 | 19.1 | 32.06 | 49.700 | 261 |
| 31 | 0.2 | 0.063 | 28.6 | 30.5 | 33.0 | 6.0 | 10.15 | 17.000 | 487 |
| 32 | 0.1 | 0.035 | 27.2 | 28.6 | 31.2 | 6.0 | 10.15 | 17.000 | 703 |
| 33 | 1.1 | 0.089 | 28.5 | 31.0 | 33.0 | 6.4 | 12.81 | 22.300 | 504 |
| 34 | 0.2 | 0.053 | 29.2 | 31.5 | 33.3 | 1.5 | 7.84 | 15.400 | 262 |
| 35 | 0.3 | 0.056 | 29.3 | 30.9 | 32.2 | 6.3 | 14.22 | 23.900 | 238 |
| 36 | 0.5 | 0.075 | 28.0 | 30.6 | 32.6 | 6.3 | 14.22 | 23.900 | 774 |
| 37 | 0.7 | 0.061 | 29.9 | 31.9 | 35.2 | 5.0 | 9.24 | 12.000 | 487 |

**ADDITIONAL INFORMATION 1 - HISTOGRAMS REPRESENTING VARIABLES DISTRIBUTION AMONG SAMPLING SITES**

Histograms representing the distribution of the variables included in the regression models.

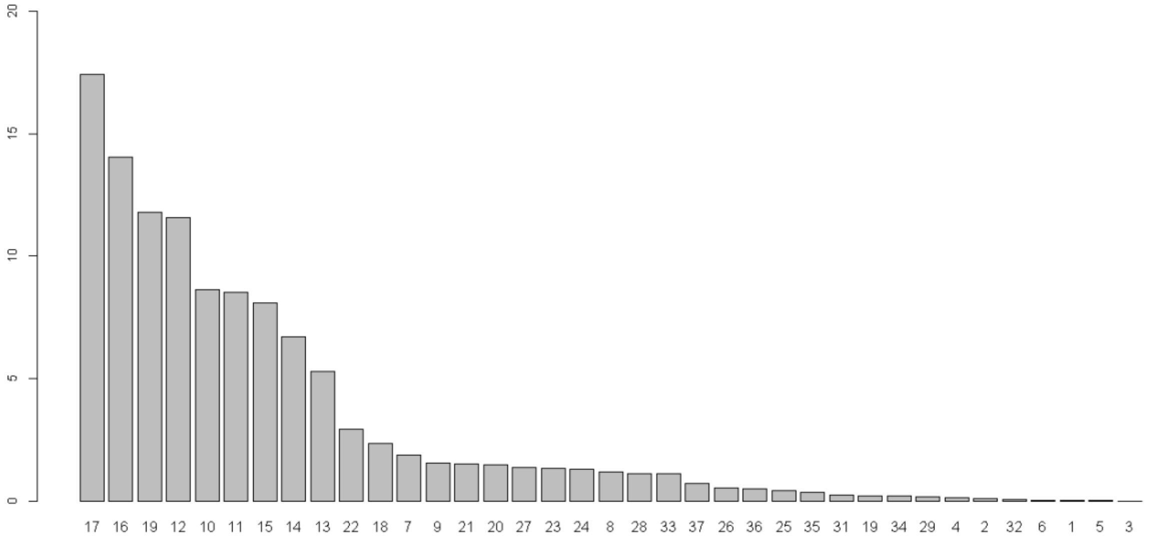


Impervious/semi natural ratio along the sampling sites of the urbanization gradient (range 0 -17.4)

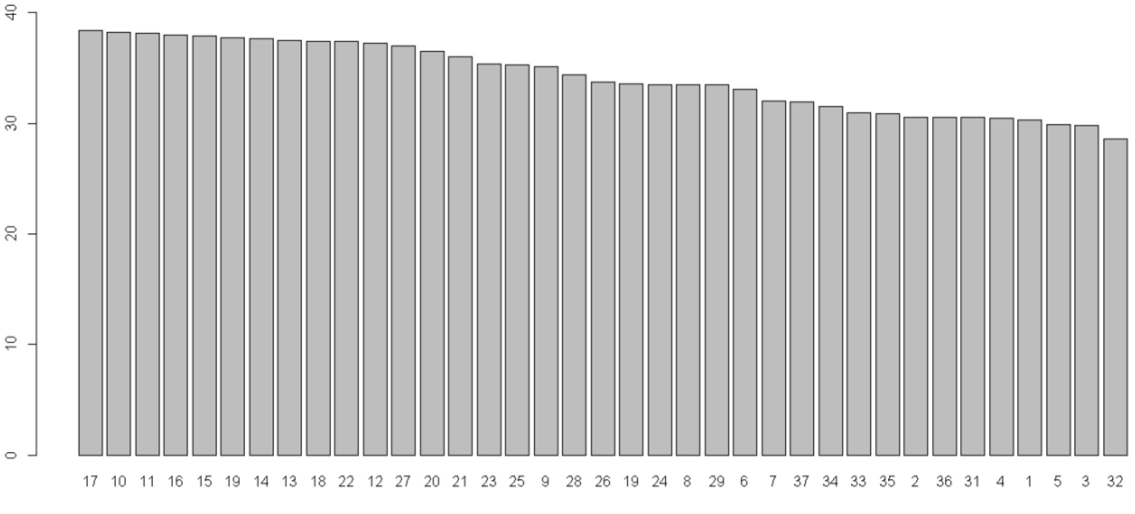


Distribution of temperature (C°) values along the sampling sites of the urbanization gradient (mean range 28.6° – 38.4°)

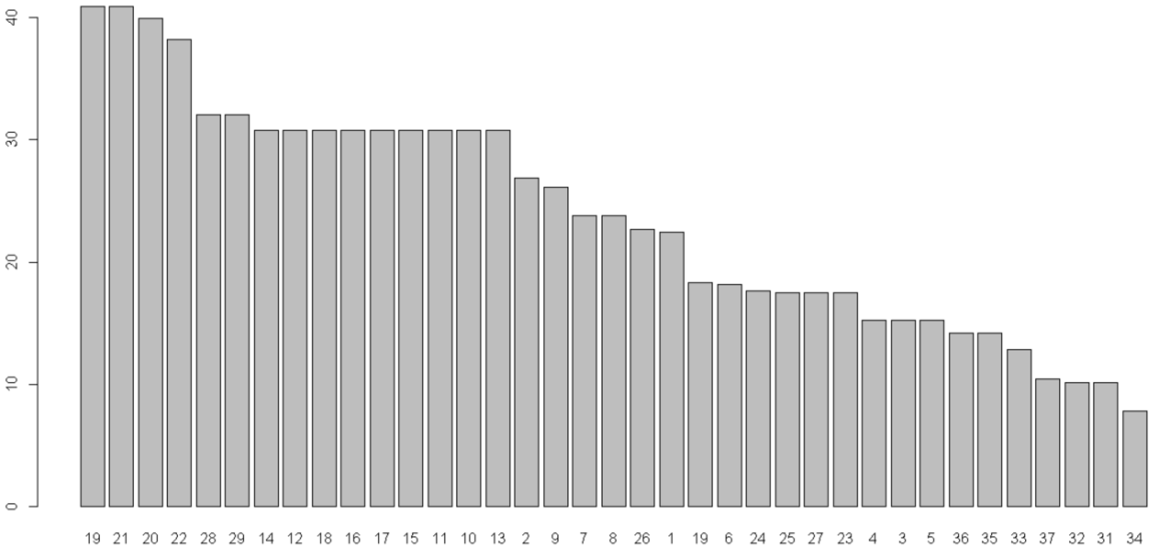


Distribution of NO2 (µg/m$^3$) values along the sampling sites constituting the urbanization gradient (mean range 23.7 µg/m3 -40.8 µg/m$^3$)

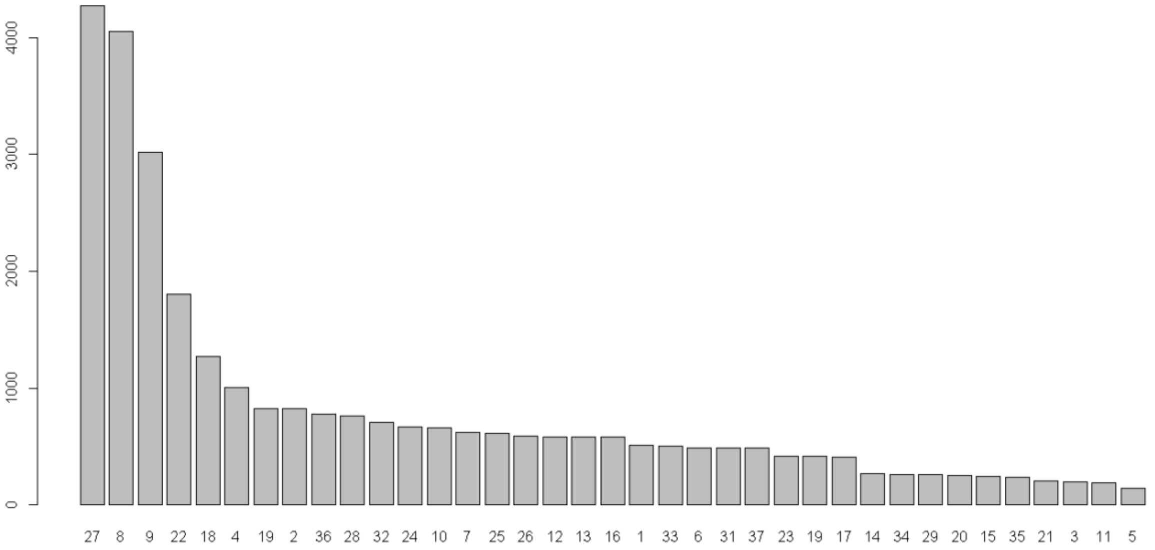


Distribution of Floral resource availability (estimated number of flowers in standardized units of space) along sampling sites constituting the urbanization gradient (range 141- 4270)

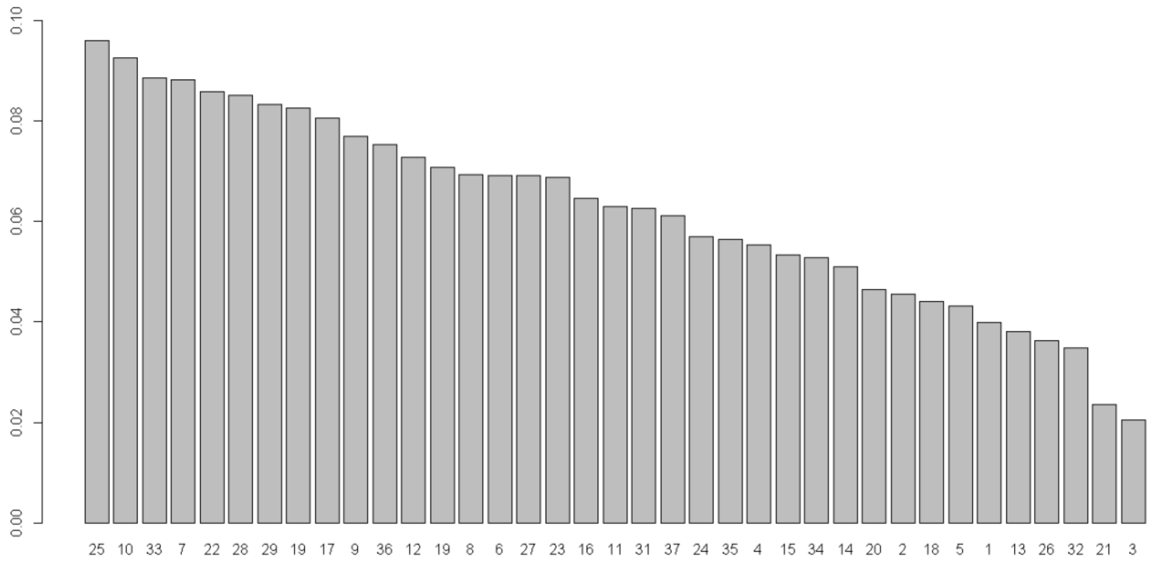


Distribution of Edge density values (measure of green patches fragmentation) along the sampling sites constituting the urbanization gradient

### TABLE S2 - Correlation matrix between variables

| | NO2 | Edge density | Temperature (C°) | Impervious/natural surfaces | Floral resources |
|---|---|---|---|---|---|
| NO2 | **1.00** | **0.09** | **0.75** | **0.64** | **0.02** |
| Edge density | **0.09** | **1.00** | **0.27** | **0.28** | **0.35** |
| Temperature (C°) | **0.75** | **0.27** | **1.00** | **0.84** | **0.12** |
| Impervious/natural surfaces | **0.64** | **0.28** | **0.84** | **1.00** | **0.03** |
| Foral resources | **0.02** | **0.35** | **0.12** | **0.03** | **1.00** |

### FIGURE S1 - Map of mean temperature and $NO_2$ sampling points

The following map shows the mean temperature in the may-july period. The location from where $NO_2$ values were recovered from the same period are also reported with a triangle. Sampling site locations are pointed by dots

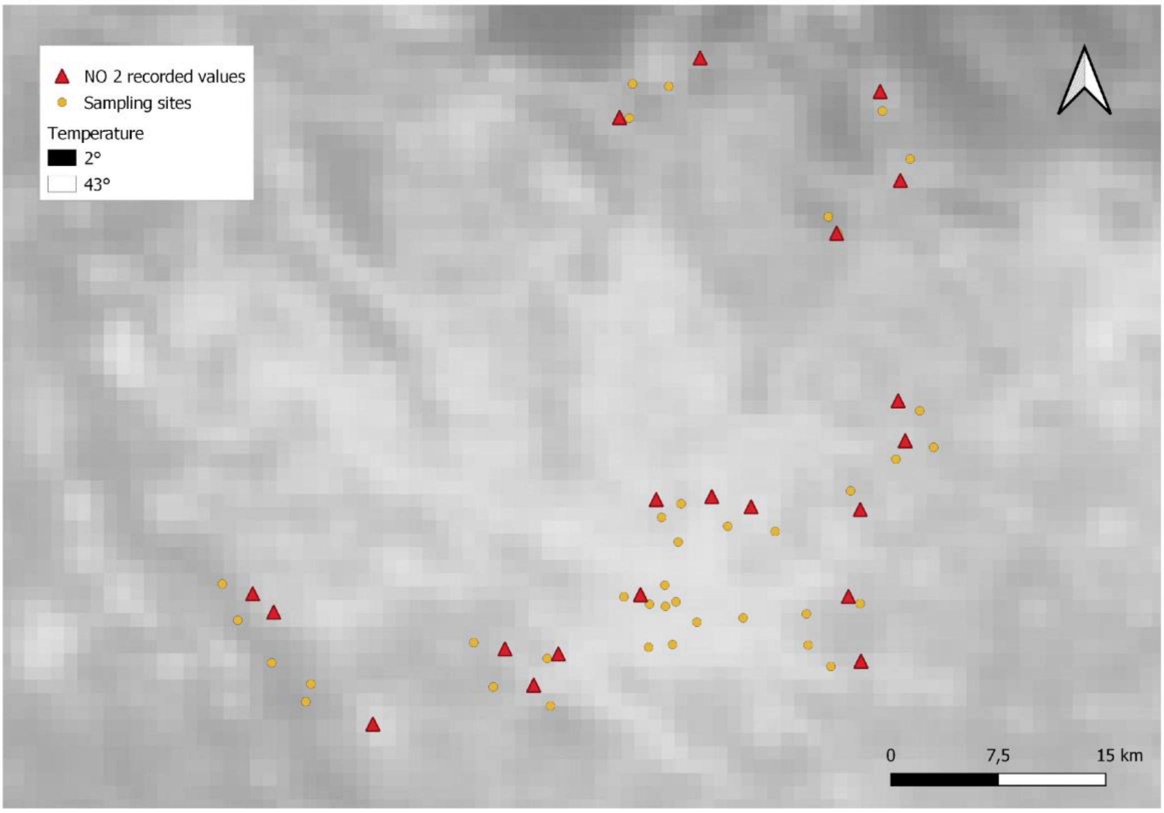

### TABLE S3 COMPLETE REGRESSION MODELS OUTPUTS

Output of Linear mixed models of body size (N= 348) and fluctuating size asymmetry (size FA) (N= 347) of each species as a function of biotic and abiotic covariates of urbanization, with site identity as random factor. Final models were selected through backward stepwise selection using AIC criterion. ΔAIC reports the difference in AIC values between full and final models. βi: regression coefficient; χ2: chi square values; df: degrees of freedom.

| Species | Response variable | Full model covariates | Final model covariates | ΔAIC | $B_i$ | χ2; df | p value |
|---|---|---|---|---|---|---|---|
| *Bombus terrestris* | Body size | Temperature | log (Floral resources) | 16.5 | 0.025 | 6.610;1 | **0.010** |
| | | Edge density | (1\|Site) | | | | |
| | | log (Floral resources) | | | | | |
| | | Variables interaction | | | | | |
| | | (1\|Site) | | | | | |
| *B.pascuorum* | Body size | Temperature | Temperature | 17.9 | -0.003 | 7.403;1 | **0.006** |
| | | Edge density | (1\|Site) | | | | |
| | | log (Floral resources) | | | | | |
| | | Variables interaction | | | | | |
| | | (1\|Site) | | | | | |
| *B.terrestris* | Size FA | Temperature | Temperature | 23.5 | 0.052 | 7.183 | **0.007** |
| | | log (Floral resources) | (1\|Site) | | | | |
| | | $NO_2$ | | | | | |
| | | Variables interaction | | | | | |
| | | (1\|Site) | | | | | |
| *B.pascuorum* | Size FA | Temperature | log (Floral resources) | 30.3 | -0.161 | 6.118;1 | **0.013** |
| | | log (Floral resources) | (1\|Site) | | | | |
| | | $NO_2$ | | | | | |
| | | Variables interaction | | | | | |
| | | (1\|Site) | | | | | |
| *B.terrestris* | Shape FA | Temperature | | | | 0.819;1 | 0.366 |
| | | log (Floral resources) | | | | 1.088;1 | 0.297 |
| | | $NO_2$ | | | | 0.774;1 | 0.379 |
| | | Variables interaction | | | | 0.689;1 | 0.407 |
| | | (1\|Site) | | | | | |
| *B.pascuorum* | Shape FA | Temperature | | | | 2.508;1 | 0.113 |
| | | log (Floral resources) | | | | 0.023;1 | 0.881 |
| | | $NO_2$ | | | | 0.624;1 | 0.429 |
| | | Variables interaction | | | | 0.675;1 | 0.411 |
| | | (1\|Site) | | | | | |

### ADDITIONAL INFORMATION - 2 Description of DUSAF levels categorised as "Impervious" and "Semi Natural"

List of Level 3 and 4 DUSAF codes Categorized as "Impervious":

"111", "1121", "1122", "1123", "1211", "1212", "122", "124", "131", "132", "133", "134", "1421", "1422", "1423"

and "Seminatural": "141", "224", "231", "311", "314", "322", "324", "331", "332", "411".

Full explanation of codes is available at:

https://www.cartografia.regione.lombardia.it/metadata/Dusaf/doc/Legenda_DUSAF_2018_6_0.pdf
